# Noninvasive prenatal detection of fetal trisomy and single gene disease by shotgun sequencing of placenta originated exosome DNA: a proof-of-concept validation

**DOI:** 10.1101/464503

**Authors:** Weiting Zhang, Sen Lu, Jia Zhao, Dandan Pu, Haiping Zhang, Lin Yang, Peng Zeng, Fengxia Su, Zhichao Chen, Mei Guo, Ying Gu, Yanmei Luo, Huamei Hu, Yanping Lu, Hongyun Zhang, Fang Chen, Ya Gao

## Abstract

**Background:** During human pregnancy, Placental trophectoderm cells can release exosomes into maternal circulation. Trophoblast cells also give rise to cell-free DNA (cfDNA) and has been used for noninvasive prenatal screening for chromosomal aneuploidy. We intended to prove the existence of exosomal DNA (exoDNA) in the exosomes of maternal blood and compared exoDNA with plasma cfDNA in terms of genome distribution, fragment length, and the possibility of detecting genetic diseases.

**Methods:** Maternal blood from 20 euploid pregnancies, 9 T21 pregnancies, 3 T18 pregnancies, 1 T13 pregnancy and 2 pregnancies with FGFR3 mutations were obtained. Exosomes enriched from maternal plasma were confirmed by transmission electronic microscopy (TEM), western blotting and flow cytometry. ExoDNA was extracted and its fetal origin was confirmed by realtime fluorescence quantitative PCR(Q-PCR). Besides, exoDNA content was uncovered by Q-PCR. To characterize exoDNA and compare with cfDNA, pair-end whole genome sequencing was performed. Lastly, the fetal risk of genetic diseases was analyzed using the exoDNA sequencing data.

**Results:** ExoDNA span on all 23 pairs of chromosomes and mitochondria, sharing a similar distribution pattern and higher GC content comparing with cfDNA. ExoDNA showed shorter fragments yet lower fetal fraction than cfDNA. ExoDNA could be used to determine fetal gender correctly, and all trisomies as well as de novo FGFR3 mutations.

**Conclusions:** We proved that fetal exoDNA could be identified in the exosomes extracted from maternal plasma. ExoDNA shared some similar features to cfDNA and could potentially be used to detect genetic diseases in fetus.

## Introduction

Exosomes are extracellular vesicles of 50 to 100 nanometer in diameter, and covered by lipid bilayer membrane carrying tetraspanins markers such as CD9 and CD63, and CD81 [1]. Exosomes, acting as an intercellular communicator in different body fluid, contain nucleic acids, proteins, and lipids [1-4]. During human pregnancy, placental trophoblasts can release exosomes into maternal circulation, and the exosome concentration positively correlates with gestation weeks [5,6]. Although the exact role of placental exosomes is still unclear, the concentration and composition balance of exosome may be important in regulating some key pregnancy activities, such as immune tolerance, maternal-fetal surface remodeling, and inflammation response [7,8]. For instance, Saloman *et al*. found that patients with gestational diabetes had altered concentration and bioactivity of placenta-derived exosomes [9]. Luo *et al*. reported that human villous trophoblasts secreted placenta-specific microRNA into maternal plasma via exosomes, and may confer antiviral ability though communicating with target cells [10]. Baig *et al*. showed that in the blood of pregnant women with preeclampsia, there was an elevated concentration of syncytiotrophoblast microvesicles, and the proteomic and lipidomic profiles of the microvesicles were considerably changed comparing with normal pregnant women [11,12].

Recently, several studies demonstrated that tumor derived exosomes contain exosomal DNA (exoDNA). For instance, in cancer cell lines and serum from patients with pancreatic cancer, tumor-derived exosomes contained double-stranded DNA fragments distributing on the entire genome, which could be used to detect tumor related genes [13-15]. Similarly, in exosomes isolated from visceral cancers, exoDNA has been used to detect copy number variation, point mutations and gene fusions by the whole genome and whole exome sequencing[16]. Mitochondrial DNA (mtDNA) has also been identified in exosomes from astrocytes and glioblastoma cells [17]. However, the evidence of placenta derived exoDNA in the maternal circulation during pregnancy has rarely been reported. In this study, we intended to provide the evidence that exoDNA exists in placental exosomes in maternal circulation.

Placenta trophoblast cells are known to release cell-free DNA (cfDNA) into maternal circulation, which are DNA fragments of 160-170 base pair [18]. The concentration of placental cfDNA in plasma, also known as the fetal fraction, is on average 10% of the total plasma cfDNA, and increases with gestational weeks [19]. Recently, plasma cfDNA is used to noninvasively detect fetal conditions such as chromosome aneuploidy and CNV [20-22]. If exoDNA exists in placental exosomes and enters maternal circulation, it would be interesting to compare the molecular features of exoDNA with plasma cfDNA. Therefore, we characterized the genome distribution, fragment length, and fetal fraction of exoDNA using massively parallel sequencing (MPS) and compared with cfDNA. Lastly, we explored the diagnostic potential of exoDNA in prenatally detecting chromosomal abnormality and de novo mutations in fetuses.

## Result

### Determining the presence of placenta derived exosomes in maternal plasma

Exosomes from 500ul plasma of 20 normal pregnancies (Table S1) were isolated using a commercial kit and visualized by TEM. Figure 1A shows the bilayer-membrane exosomes with the typical cap-shape looking and the size of 30nm-50nm in diameter. Western blotting analysis of typical membrane markers of exosome showed the presence of CD9, CD63 and CD81 (Figure 1B). This was confirmed by flow cytometry analysis of CD9 and CD63 expression of exosomes (Figure 1C). We also used the Western blotting to confirm the expression of placental alkaline phosphatase (PLAP) in exosomes extracted from maternal plasma (Figure 1B), which is a syncytiotrophoblast-specific proteins[6]. Therefore, exosomes could be isolated from the plasma of pregnant women, and placenta derived exosomes expressing the PLAP marker could be identified.

**Fig.1.**
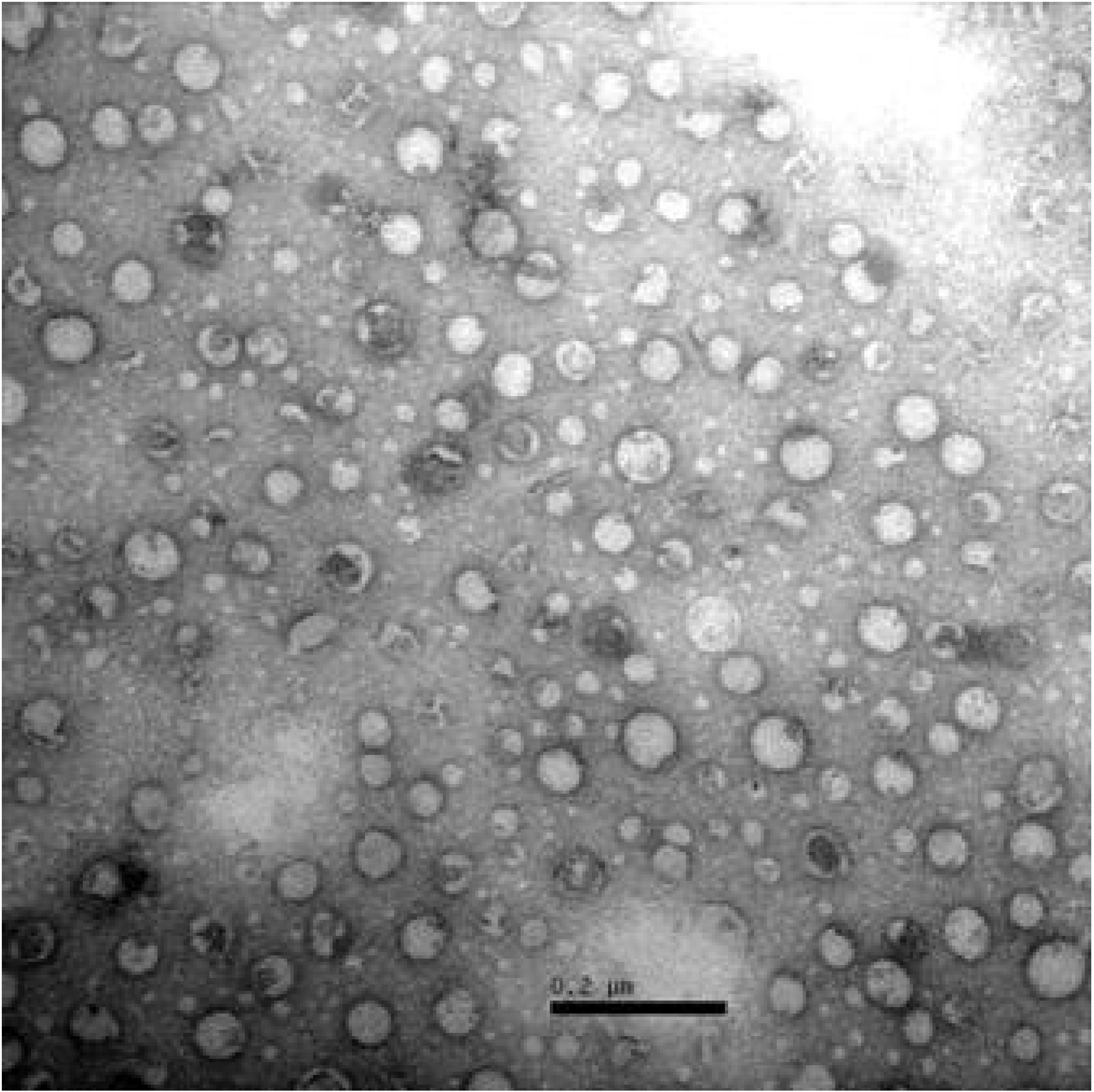
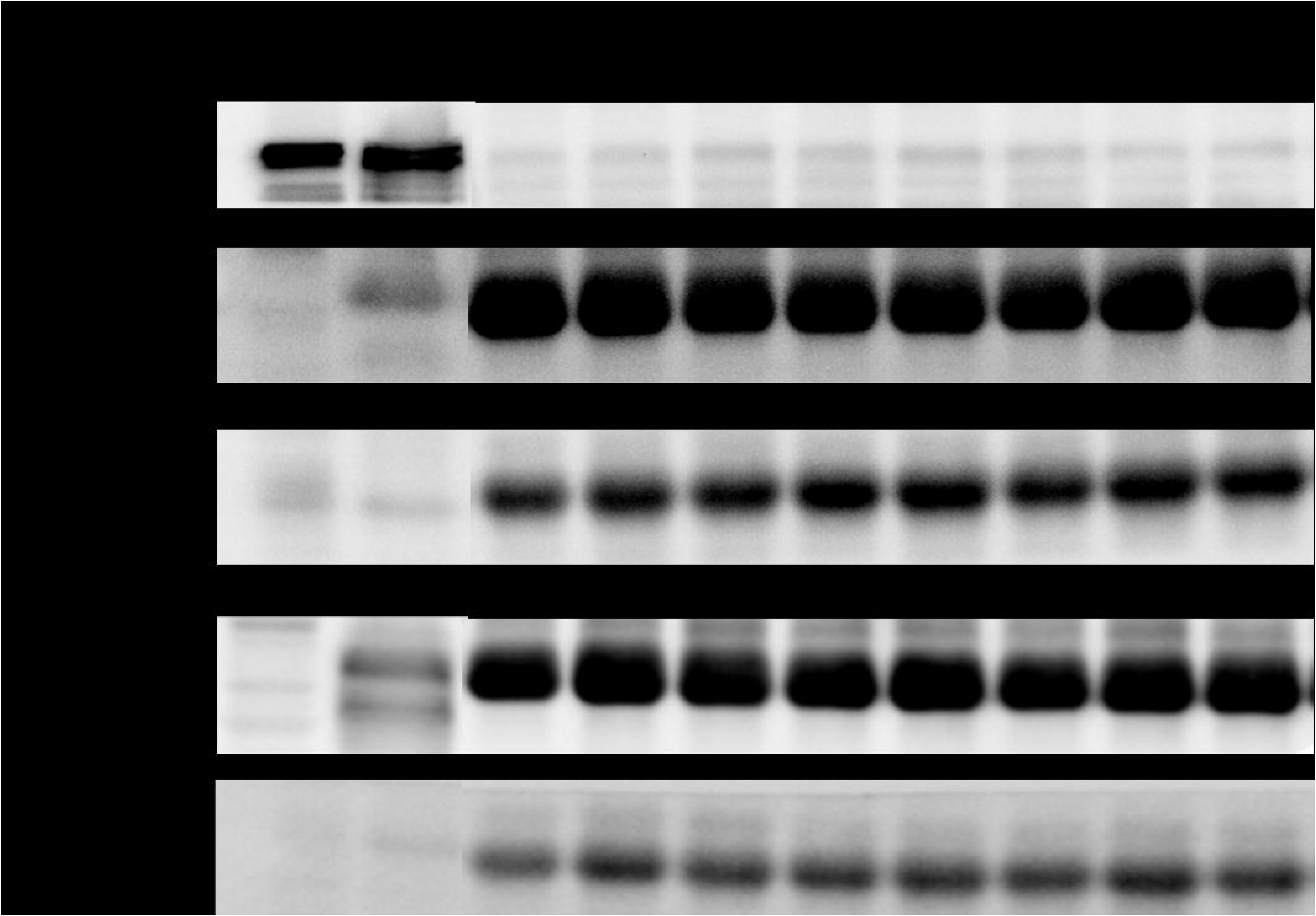
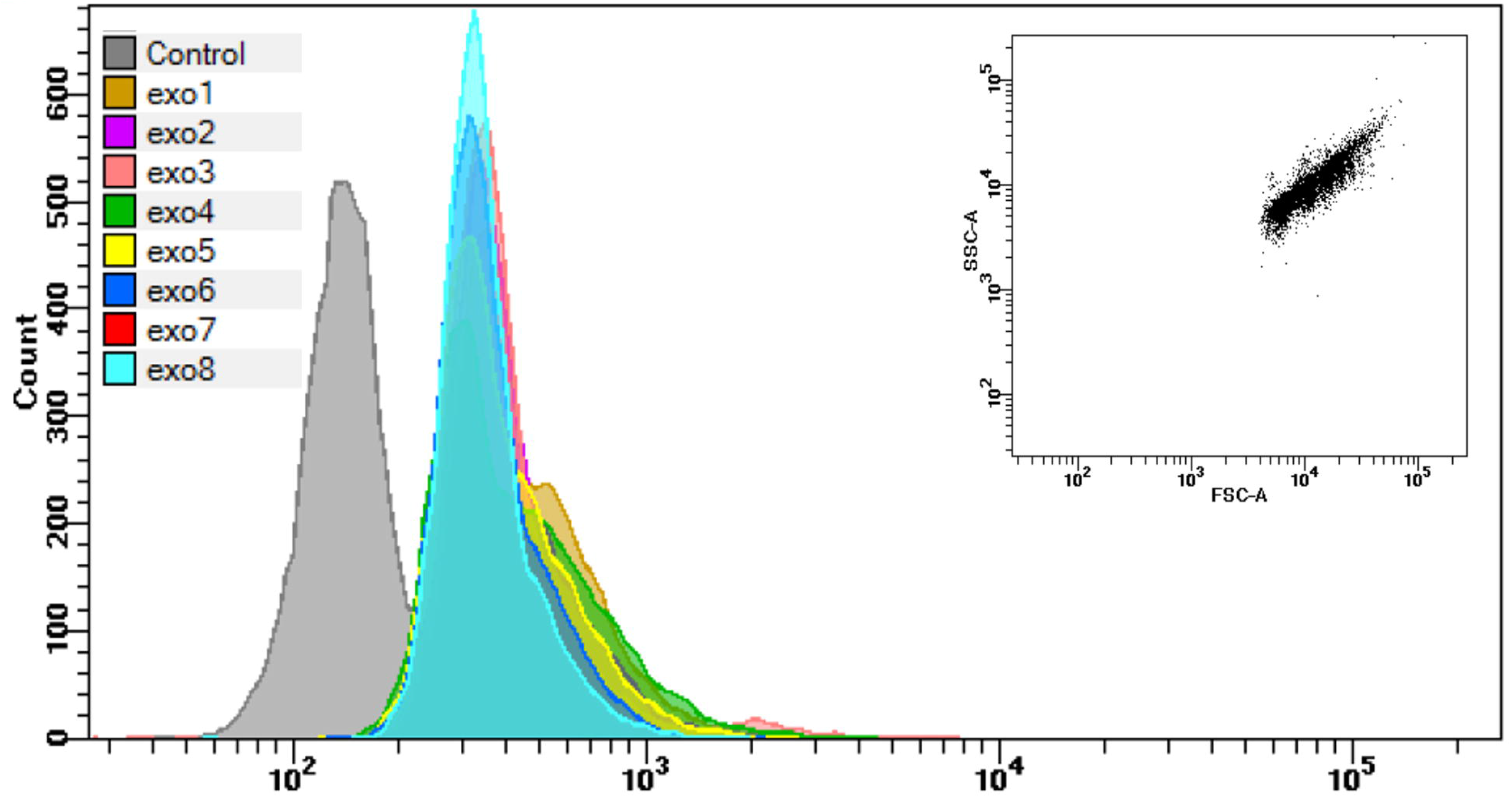
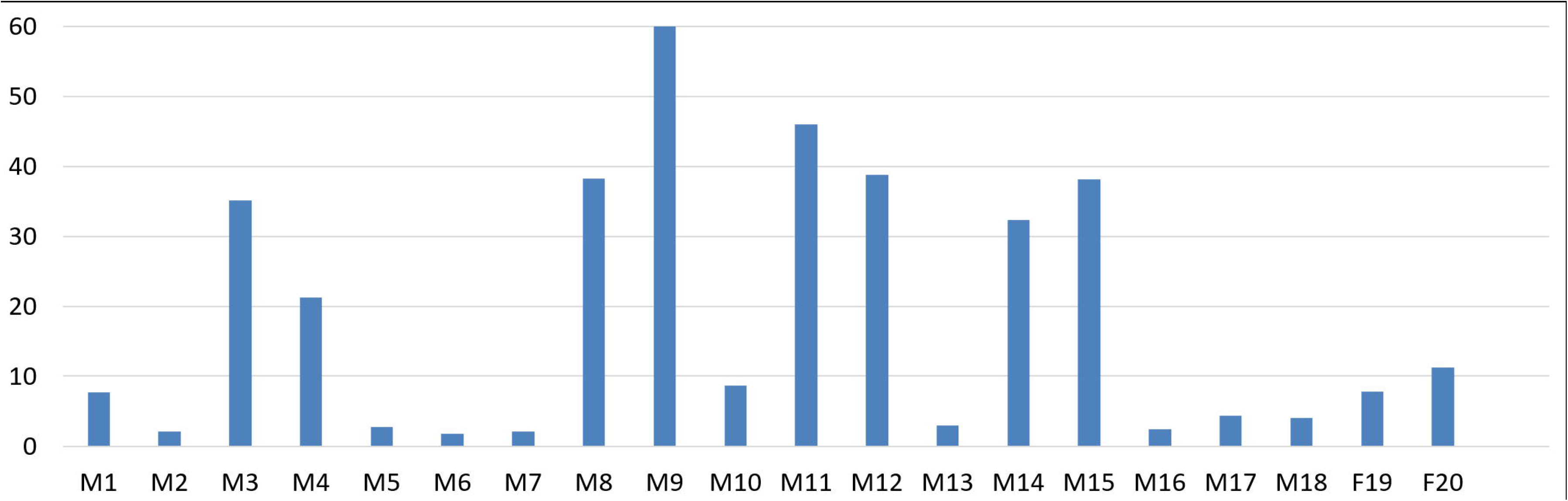
Characterization of exosomes from plasma. **(A)** transmission electron microscope detection of exosomes isolated by SBI ExoQuick kit. Scale bar=200nm; **(B)** Western blot analysis for the classical biomarker of exosomes (CD9, CD63 and CD81), placenta specific biomarker (PLAP). Endoplasmic reticulum marker calnexin was used as a control marker, exo1-8 is from eight maternal plasma samples; **(C)** flow cytometry detection for exosomes from eight maternal plasma samples, exosomes isolated with CD9 magnetic beads were stained with CD63-Alexa647 and isotype control IgG-Alexa647. **(D)** The ratio of cfDNA to exoDNA from equal volume plasma(250μl) was calculated according to the ∆Ct value. **(E)** Q-PCR of exoDNA from two maternal plasma with male fetus and female fetus respectively, long autosome DNA and short autosome DNA signal show the existence of template while Y chromosome DNA signal shows fetal gender.

### Determining the amount of exoDNA and fetal origin of exoDNA

We firstly treated the exosomes with DNaseI to avoid potential cfDNA contamination. Then DNase I was deactivated, and exoDNA was extracted to compare with the cfDNA extracted from the isochoric plasma of the same pregnant woman. We described the content of extracted cfDNA and exoDNA in elution buffer by Qubit^TM^ dsDNA HS Assay kit(Figure S1). Then, Q-PCR was conducted in both exoDNA and cfDNA using primers targeting autosomes and Y chromosome. The relative amount of exoDNA was determined by the △Ct value between cfDNA and exoDNA, which showed that the amount of exoDNA is fewer than cfDNA, ranging from 1/2.8 to 1/61.5 (Figure 1D). Nonetheless, autosomal signals in all 20 pregnancies could be detected by Q-PCR, which proves the presence of exoDNA. Moreover, Q-PCR successfully detected the signal of Y chromosome in 18 pregnancies with male fetuses and showed no signal of Y chromosome in 2 pregnancies with female fetuses (Figure 1E). Thus, exosomes from the plasma of pregnant women contained exoDNA of fetal origin, and fetal gender could be detected using exoDNA.

### Genome distribution of exoDNA by MPS

To characterize the molecular features of exoDNA, we constructed the DNA libraries and performed pair-end (PE) low-coverage MPS using the exoDNA obtained from the above 20 pregnant women. Plasma cfDNA from the same 20 pregnant women was sequenced as comparison. We obtained on average 13.1 million(7.7M) unique reads (0.25X) of exoDNA in each sample(Table 1). Figure 2 shows the distribution of aligned PE reads of exoDNA and cfDNA on human genome. Both exoDNA and cfDNA displayed reads coverage on all 23 pairs of chromosomes. The coefficient of variation (CV) of the relative read counts of exoDNA and cfDNA in 20 samples showed no statistical difference (P=0.125), with the medium value of 0.131 and 0.149 respectively, which indicates that both DNAs evenly distributed on human genome (Figure 3A). However, the GC content of exoDNA was constantly 1.16 times higher than that of cfDNA (Table 1 and Figure 3B). Moreover, mitochondrial DNA (mtDNA) was detected in exosome by MPS, and the reads percentage of mitochondrial exoDNA was on average 2.18 times higher than that of cfDNA (p<0.001) (Table 1 and Figure 3C).

**Fig.2.**
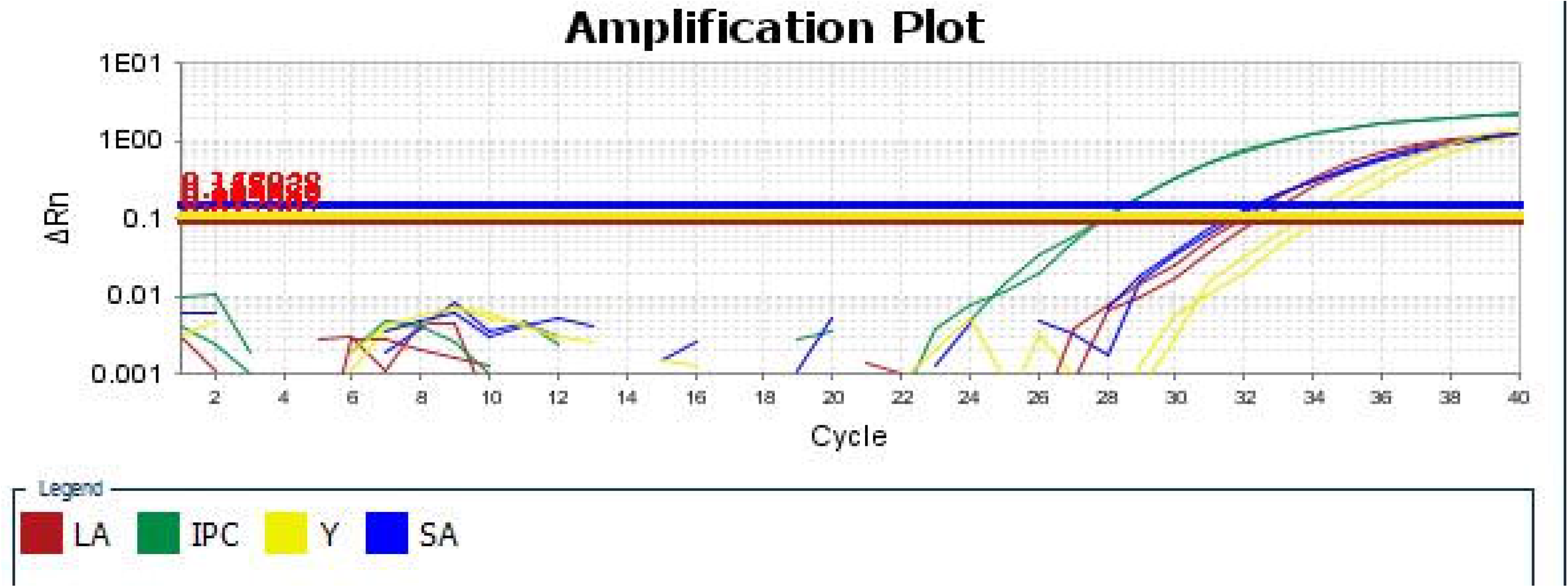
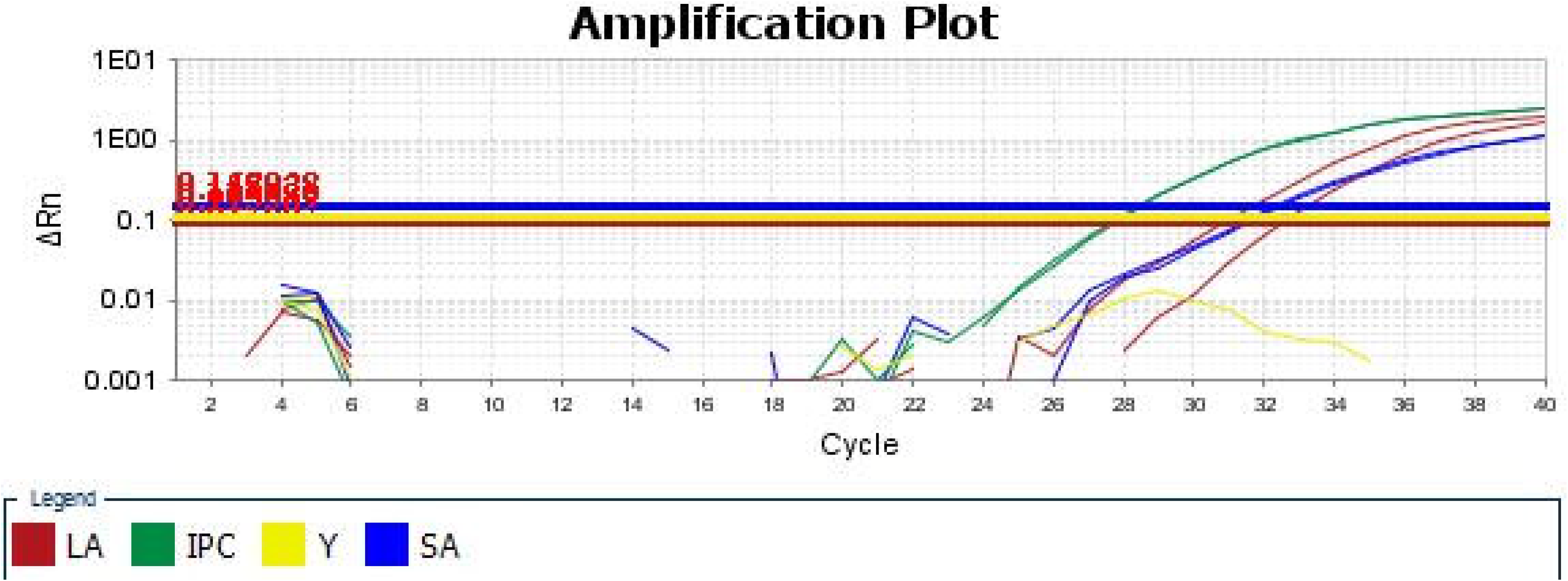
Whole genome distribution of exoDNA(blue) and cfDNA(red) of sample exo16 illustrated by Circos(outer to inner: reads ratio; GC content);

**Table 1.** Sequencing data of the 20 Euploidy.

| Sample ID |  | Effective reads/M | Depth | CV mean | GC% | mtDNA% | cffDNA% |
| --- | --- | --- | --- | --- | --- | --- | --- |
| Euploidy-1 | exo | 9.80 | 0.32X | 0.1006 | 46.99 | 0.0201 | 5.48 |
|  | cfDNA | 2.57 | 0.08X | 0.1625 | 40.04 | 0.0136 | 11.90 |
| Euploidy-2 | exo | 4.93 | 0.16X | 0.1315 | 46.25 | 0.0106 | 5.48 |
|  | cfDNA | 2.49 | 0.08X | 0.1659 | 39.71 | 0.0087 | 11.90 |
| Euploidy-3 | exo | 8.85 | 0.29X | 0.0966 | 43.67 | 0.0203 | 5.84 |
|  | cfDNA | 3.63 | 0.12X | 0.1385 | 39.42 | 0.0079 | 8.86 |
| Euploidy-4 | exo | 16.13 | 0.53X | 0.0819 | 45.78 | 0.0063 | 6.66 |
|  | cfDNA | 5.23 | 0.17X | 0.1208 | 39.10 | 0.0023 | 12.76 |
| Euploidy-5 | exo | 10.36 | 0.34X | 0.1109 | 49.00 | 0.0133 | 2.74 |
|  | cfDNA | 3.59 | 0.12X | 0.1398 | 39.37 | 0.0055 | 6.08 |
| Euploidy-6 | exo | 2.42 | 0.08X | 0.1971 | 47.28 | 0.0146 | 1.27 |
|  | cfDNA | 5.24 | 0.17X | 0.1184 | 38.92 | 0.0026 | 3.76 |
| Euploidy-7 | exo | 4.59 | 0.15X | 0.1316 | 46.19 | 0.0090 | 2.94 |
|  | cfDNA | 4.67 | 0.15X | 0.1229 | 39.46 | 0.0021 | 8.74 |
| Euploidy-8 | exo | 9.57 | 0.31X | 0.1055 | 46.75 | 0.0082 | 2.06 |
|  | cfDNA | 14.89 | 0.48X | 0.0755 | 39.70 | 0.0020 | 5.64 |
| Euploidy-9 | exo | 5.93 | 0.19X | 0.1297 | 48.12 | 0.0081 | 1.60 |
|  | cfDNA | 10.11 | 0.33X | 0.0951 | 40.62 | 0.0030 | 5.64 |
| Euploidy-10 | exo | 2.37 | 0.08X | 0.1804 | 46.68 | 0.0080 | 1.71 |
|  | cfDNA | 4.54 | 0.15X | 0.1234 | 40.14 | 0.0061 | 7.90 |
| Euploidy-11 | exo | 2.60 | 0.08X | 0.1704 | 45.71 | 0.0094 | 4.38 |
|  | cfDNA | 2.03 | 0.07X | 0.1814 | 39.16 | 0.0040 | 11.14 |
| Euploidy-12 | exo | 17.03 | 0.55X | 0.0854 | 47.29 | 0.0049 | 2.92 |
|  | cfDNA | 3.11 | 0.10X | 0.1502 | 38.82 | 0.0020 | 5.18 |
| Euploidy-13 | exo | 7.41 | 0.24X | 0.1166 | 47.04 | 0.0138 | 1.86 |
|  | cfDNA | 2.39 | 0.08X | 0.1672 | 40.18 | 0.0029 | 6.38 |
| Euploidy-14 | exo | 5.63 | 0.18X | 0.1279 | 46.96 | 0.0107 | 1.53 |
|  | cfDNA | 1.77 | 0.06X | 0.1941 | 39.03 | 0.0066 | 3.94 |
| Euploidy-15 | exo | 29.38 | 0.96X | 0.0621 | 44.67 | 0.0097 | 8.62 |
|  | cfDNA | 3.47 | 0.11X | 0.1417 | 38.43 | 0.0032 | 13.16 |
| Euploidy-16 | exo | 2.93 | 0.10X | 0.1543 | 42.26 | 0.0227 | 7.24 |
|  | cfDNA | 1.76 | 0.06X | 0.1932 | 39.51 | 0.0141 | 9.00 |
| Euploidy-17 | exo | 1.99 | 0.06X | 0.1830 | 44.37 | 0.0122 | 4.90 |
|  | cfDNA | 1.67 | 0.05X | 0.1989 | 38.97 | 0.0028 | 5.94 |
| Euploidy-18 | exo | 6.58 | 0.21X | 0.1055 | 41.90 | 0.0296 | 8.24 |
|  | cfDNA | 3.02 | 0.10X | 0.1506 | 39.64 | 0.0166 | 9.74 |
| Euploidy-19 | exo | 4.30 | 0.14X | 0.1364 | 45.03 | 0.0068 | NA |
|  | cfDNA | 1.54 | 0.05X | 0.2073 | 38.76 | 0.0034 | NA |
| Euploidy-20 | exo | 1.54 | 0.05X | 0.2152 | 44.37 | 0.0225 | NA |
|  | cfDNA | 4.24 | 0.14X | 0.1335 | 38.96 | 0.0098 | NA |

Notes: Abbreviations: CV : Coefficient of Variance
mtDNA : Mitochondrial DNA
cffDNA : cell free fetal DNA

**Fig.3.**
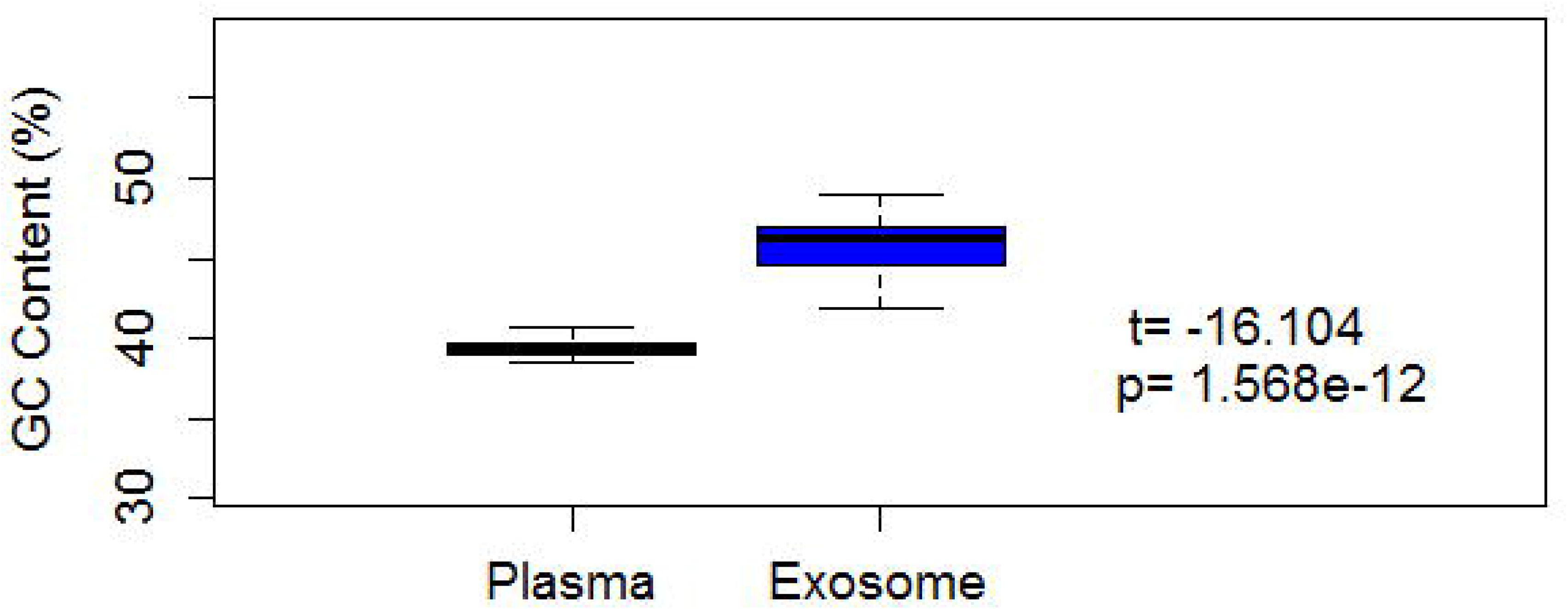
Properties of exoDNA compare with cfDNA. Coefficient of variation **(A)**, GC content **(B)**, mtDNA rate **(C)**, fragment size **(D)** and fetal fraction **(E)** of all 20 samples of cfDNA and exoDNA;

### Comparison of fragment length and fetal fraction of exoDNA and cfDNA

With the paired-end sequencing reads, we compared the DNA fragment length of exoDNA and cfDNA. Among 20 samples, cfDNA showed the median fragment size of 168.5bp with the standard deviation of ±1.25bp, as expected. In contrast, exoDNA showed shorter fragments and a more dispersive distribution (p<0.001). The median fragment size of exoDNA was 152.4bp with the standard deviation of ±10.51bp (Figure 3D and Figure S2).

We then compared the fetal fraction of exoDNA and cfDNA calculated using the chrY approach. Although the fetal fraction of exoDNA had a nice correlation to that of cfDNA (R=0.744, p<0.001), it was significantly lower than cfDNA (averagely 0.52-fold fetal fraction of cfDNA, p<0.001) and with weak relationship to gestational weeks (R=0.334, p=0.175) (Figure 3E and Figure S3).

### Detection of fetal trisomies and monogenic diseases by exoDNA sequencing

Since exoDNA demonstrated some similar characteristics and fetal origin to cfDNA, we explored the feasibility of detecting fetal diseases using exoDNA. We obtained the plasma samples from nine T21 pregnancies, three T18 pregnancies, and one T13 pregnancy (Table S1). The 20 normal pregnancies were used as controls. Exosomes enrichment and exoDNA extraction were conducted as described above. For fetal trisomy testing, exoDNA was prepared for low-coverage whole genome sequencing and analyzed using an algorithm based on binary hypothesis T-test and logarithmic likelihood ratio Z-score (see Methods). An average of 11.8 million unique reads were obtained for each sample. In the end, all positive samples of T21, T18 and T13 showed Z-scores above 3, and thus were successfully classified as trisomy high risk (Figure 4A-C). In contrast, none of the 20 normal pregnancies showed Z-scores above 3 and were classified as low risk.

**Fig.4.**
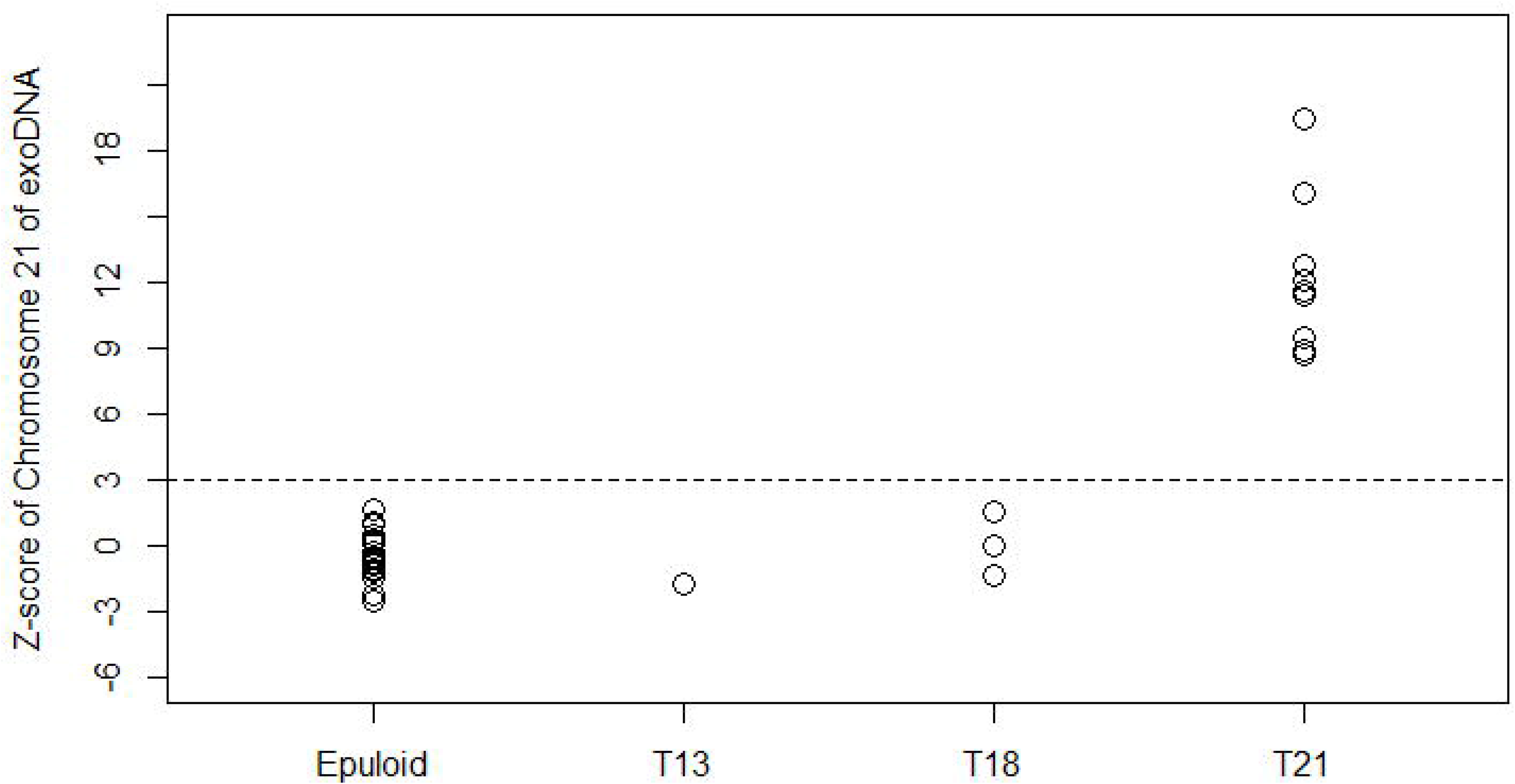
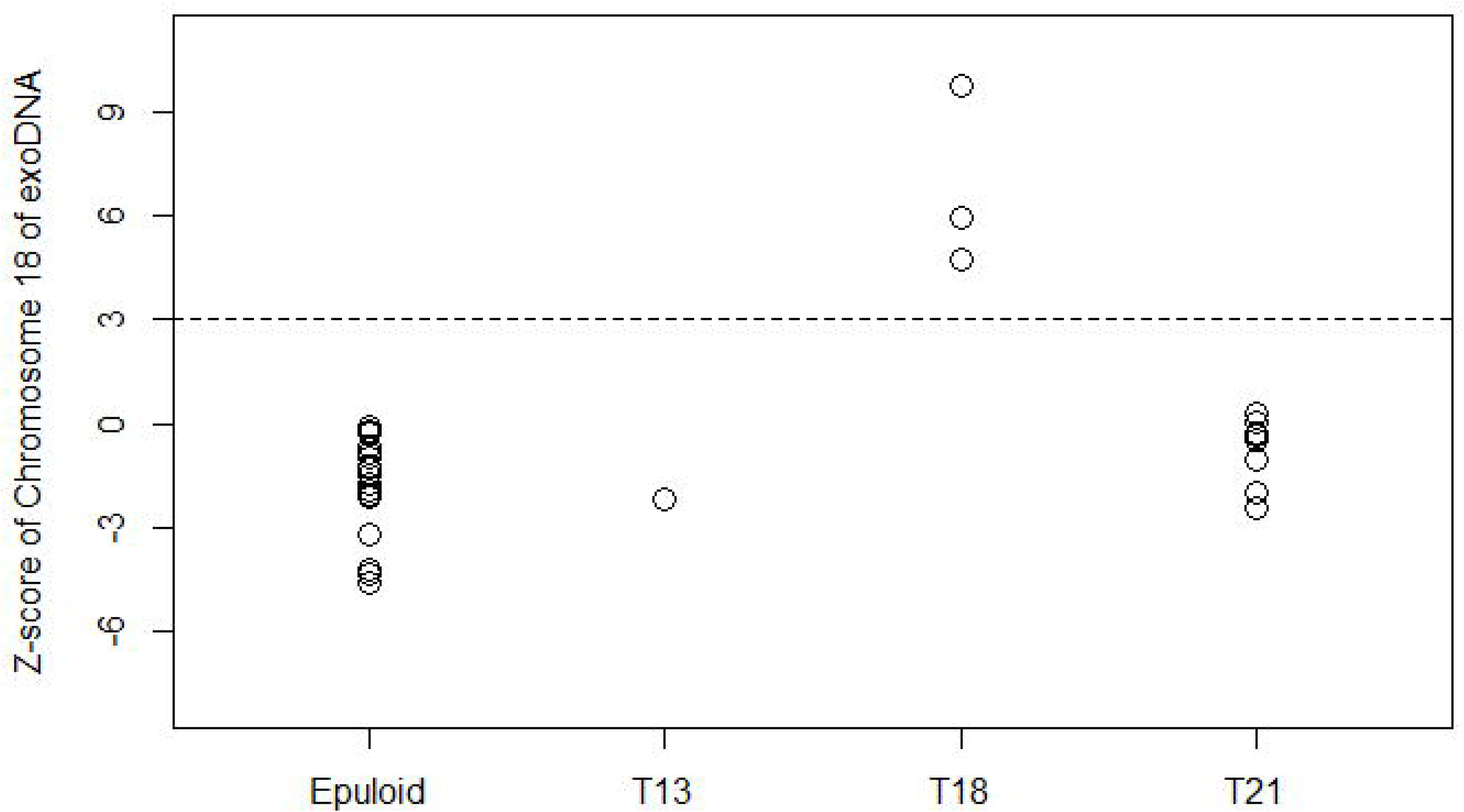
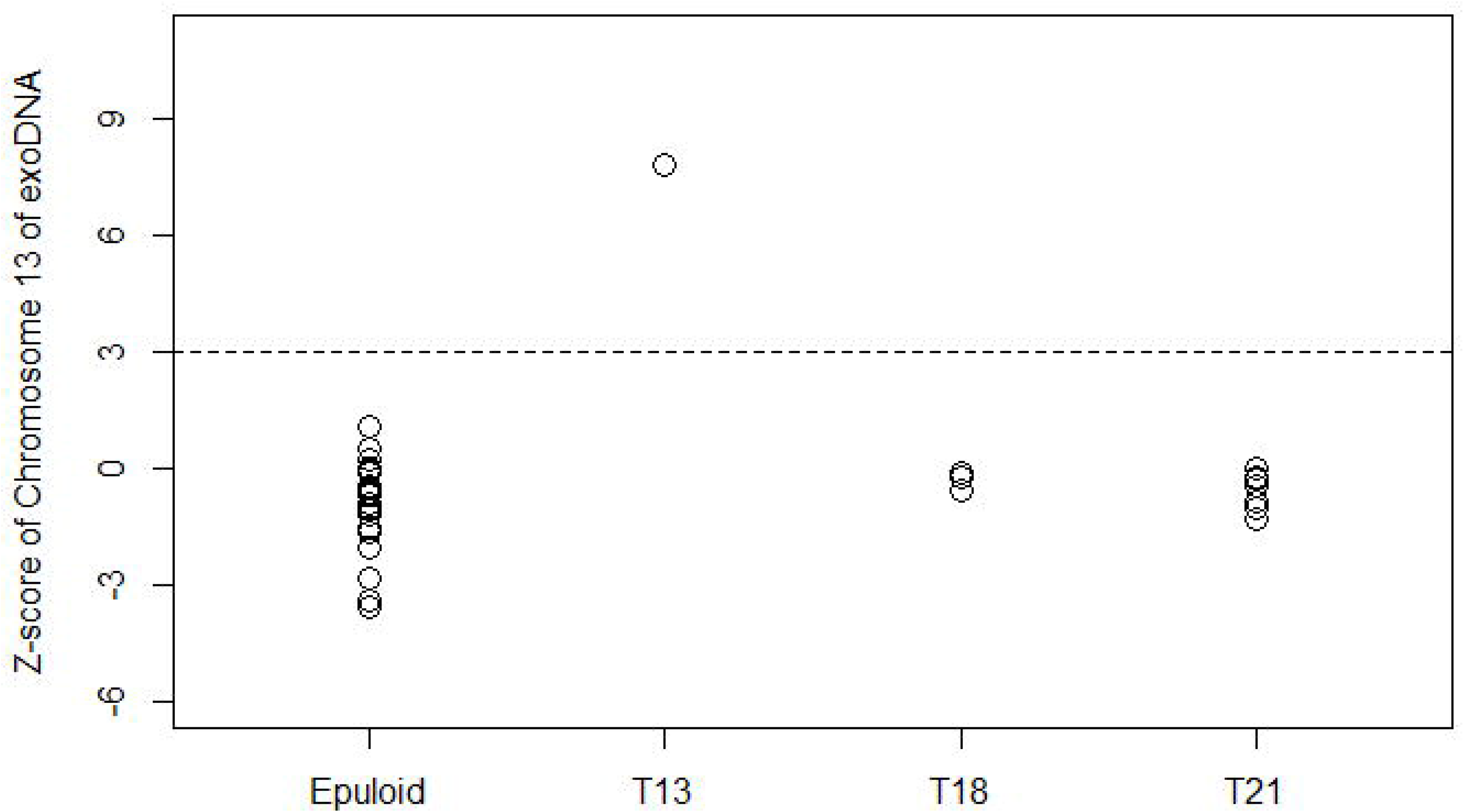
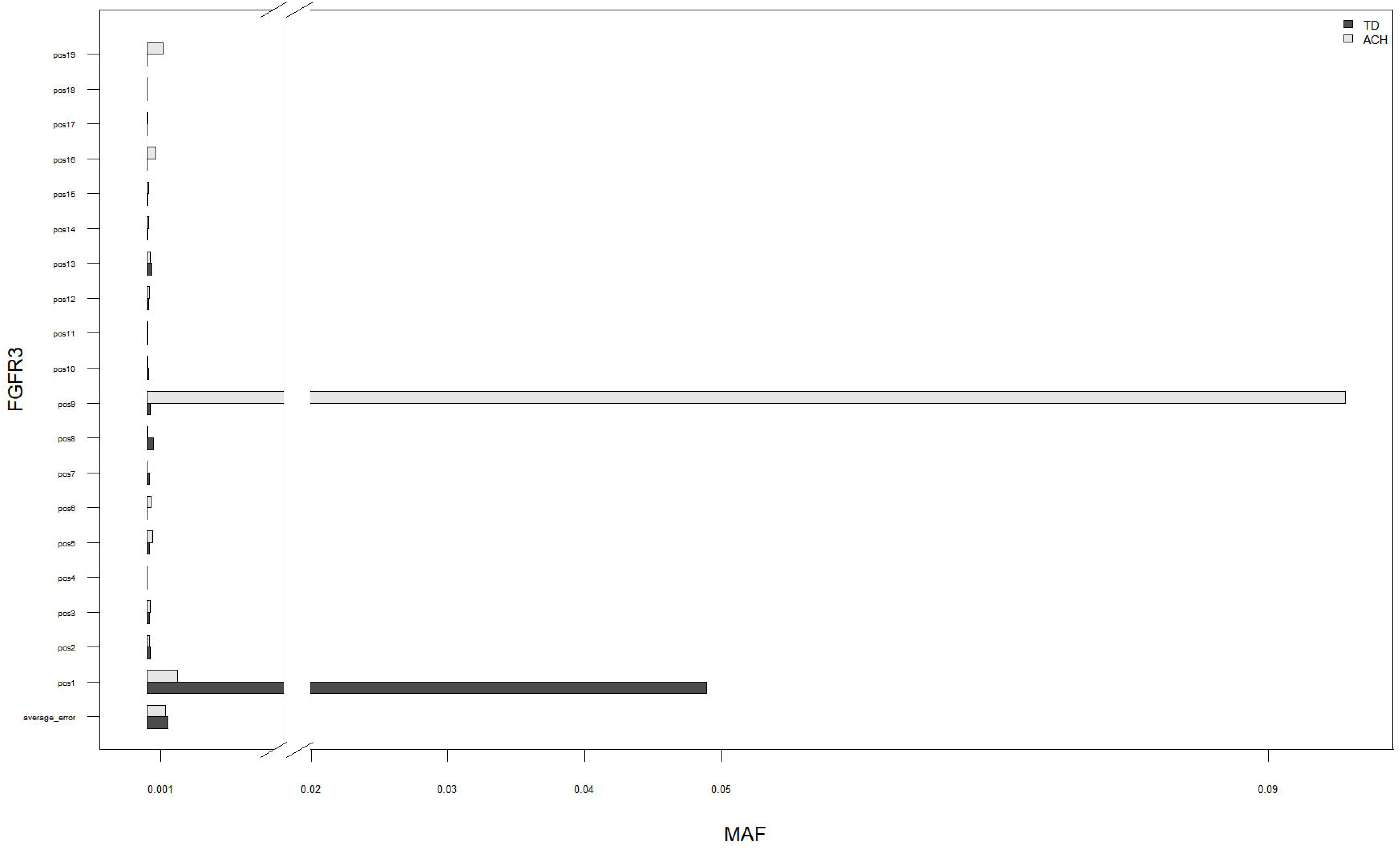
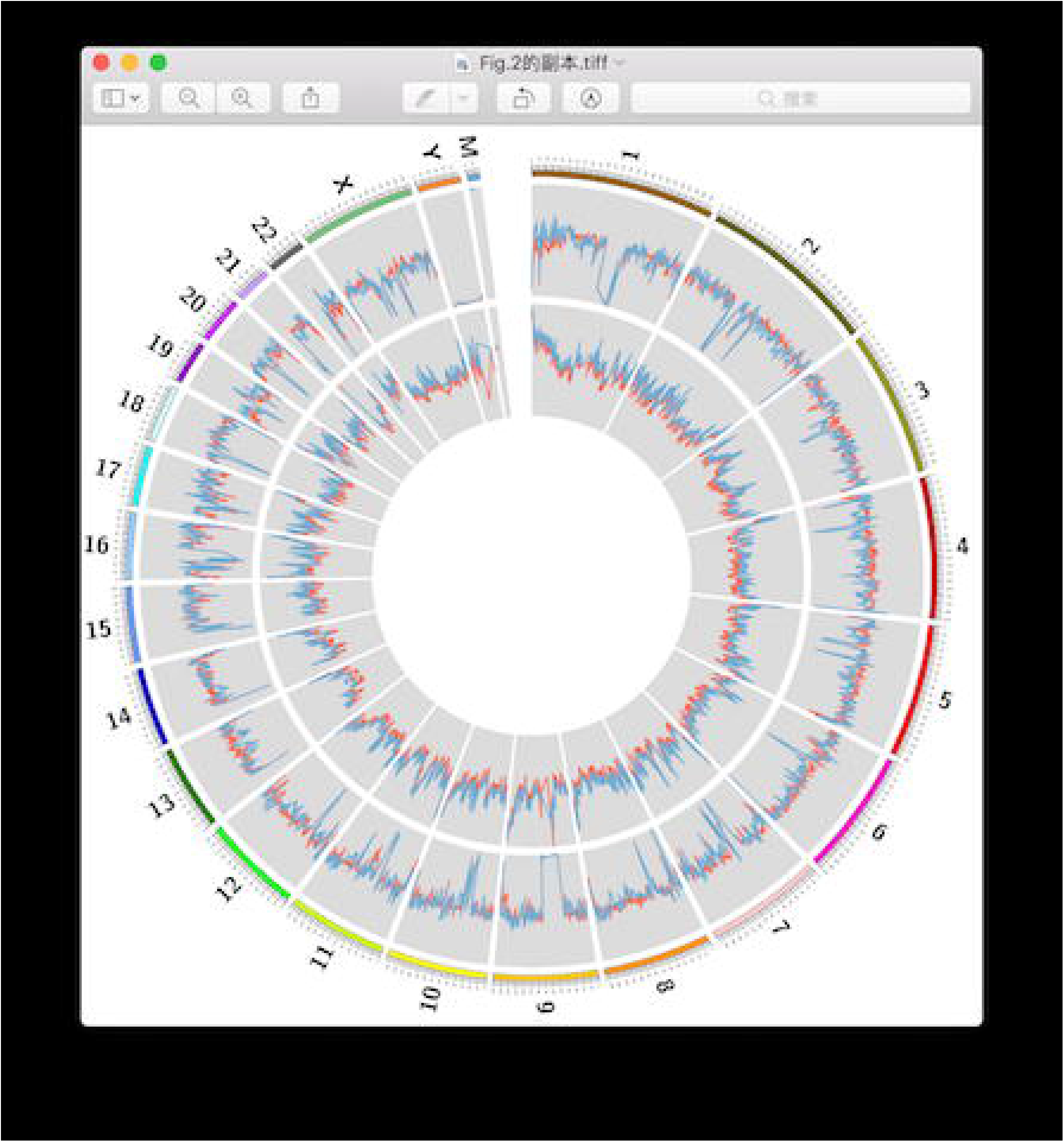
Application in diagnose of fetal diseases. z-score of exoDNA for chromosome 21**(A)**, chromosome 18**(B)**, chromosome 13**(C)** for 20 euploid, 9 trisomy 21, 3 trisomy 18 and 1 trisomy 13; (D) MAF of two mutation sites detected on FGFR3 in ACH and TD;

We further detected *de novo* mutation of monogenetic diseases using the exoDNA extracted from two pregnancies with autosomal dominant disease achondroplasia (ACH) and thanatophoric dysplasia (TD) (Table S2). Multiplex PCR was performed to amplify the coding region of *FGFR3* gene, and the amplification products were sequenced by MPS. In the test of ACH and TD, the *de novo* mutation of c.1138G>A and c.742C>T showed the MAF of 9.6% and 4.7% in the two cases respectively, which was classified as positive results (Figure 4D). All the results were confirmed by amniocentesis.

## Discussion

Previous studies have showed that human trophoblast specific marker PLAP can be detected on exosomes isolated from cell medium [23], placental explant cultures [8,24] and placental perfusate [25], indicating the presence of placenta originated exosomes in maternal circulation. *In vivo* study also demonstrated that the number of PLAP^+^ exosomes significantly increases in maternal circulation during pregnancy [5,6]. In this study, we confirmed the presence of placenta derived exosomes in the blood of pregnant women, and further demonstrated the existence of double stranded DNA in the exosomes isolated from maternal plasma. Importantly, we successfully detected fetal genders and calculate fetal fraction with exoDNA using Q-PCR and MPS experiments, indicating that exosomes isolated from maternal plasma contain not only maternal but also fetal exoDNA. To our knowledge, this is the first study that MPS was used to characterize the exoDNA inside the exosomes from maternal plasma.

By using the low-coverage MPS (~0.25X), we showed that exoDNA shares similarity of some key features to plasma cfDNA, such as the fetal origin and even distribution on human genome. We also found that like cfDNA, exoDNAs were short and double-stranded fragments with minor peaks of 10bp periodicity, which strongly indicates the existence of nucleosome structure [26]. Thus, exoDNA may be an important contributor to plasma cfDNA. As a supporting evidence, Rohan Fernando et al. recently exploited droplet digital PCR and confocal microscopy to show that exosomes in human blood of non-pregnant donors contained double strand DNA, which accounts for the main proportion of plasma cfDNA [27]. Give the similarity between exoDNA and cfDNA, it is reasonable to speculate that exoDNA may have great clinical value in detecting fetal disease as to the application of cfDNA in NIPT. Indeed, in this study we demonstrated that by using a the sequencing and analyzing strategy similar to plasma cfDNA NIPT [28,29], fetal T21, T18 and T13 were accurately detected with exoDNA in a small sample set. Meanwhile, by using targeted sequencing method, we could also accurately detect the *de novo* mutations of ACH and TD. Therefore, it is promising to use exoDNA to prenatally detect genetic diseases in fetus.

Prenatal testing with exoDNA is likely to offer extra benefits comparing with plasma cfDNA. It has been reported that neuron-specific extracellular vesicles could be obtained from human plasma by immunoprecipitation and used for disease diagnosis of neurodegenerative diseases [30]. Hong et al. reported a immunoaffinity-based capture method to isolate CD34^+^ blast-derived exosomes in the acute myeloid leukemia patients’ peripheral blood, which were biologically active for potential diagnosis use [31]. Microfluidic platforms have been also reported to specifically capture tumor exosomes from the plasma of non-small-cell lung cancer patients and glioblastoma multiforme patients [32,33]. Theoretically, using the similar approaches, pure placenta-derived exosomes can be specifically isolated with the help of placenta specific markers such as PLAP. This may raise huge benefits to develop novel NIPT methods to detect copy number variation and monogenetic disease in fetus, since the high fetal fraction of placental exoDNA can overcome the current NIPT limitations that are mainly caused by the mixture of maternal-fetal cfDNA and low fetal fraction. Another potential benefit of testing exoDNA may be the protection effect by the bilayer membrane of exosomes, and therefore confers better stability than cfDNA during blood storage and preparation. This was supported by the studies that exosomes effectively protect the mRNA/miRNA for up to 5 years from plasma and 12 months from urine [34-36].

Despite the similarity, exoDNA in the exosomes from maternal plasma also showed some distinct features to plasma cfDNA, including higher GC content, higher mtDNA percentage, shorter fragments, and lower fetal fraction. The reason behind these differences is still unknown and should be further investigated. However, a possible explanation could be the different biogenesis and turnover mechanism of exoDNA comparing with cfDNA. This also helps explain the puzzle that the exoDNA quantity in our study was only 1/2 to 1/60 of cfDNA, whereas previous studies demonstrated 13.2~100 folds increase of the number of placental exosomes during pregnancy [5,6,37]. It has been previously suggested that exoDNA might be selectively packaged into exosomes to travel between cells to induce the alteration of mRNA and protein expression [10,14,16]. A recent study proposed that cell-free telomere DNA could be packaged into exosomes, and lead to parturition after transferred to maternal circulation [38]. Another research reported that exosomes maintain cellular homeostasis by excreting harmful DNA from cells, and the inhibition of exosomes secretion led to DNA accumulation in the cytoplasma and caused the DNA damage response [39]. Thus, future study is worth investigating if placental exoDNA exerts cellular functions via exosomes and passes through the maternal-fetal barrier to enter the maternal circulation.

The current study was conducted with a limited number of samples. This may explain why the fetal fraction calculated with exoDNA did not show strong association to gestational weeks. However, even with a small sample size, this proof-of-concept study clearly demonstrated the basic features and diagnostic potential of exoDNA. In summary, this is the first study that MPS is used to prove the existence and characterize the clinical value of exoDNA in the exosomes in maternal plasma. Further studies with bigger sample size and in-depth sequencing strategy is warranty to reveal more information of the biogenesis and turnover of exoDNA.

## Materials and Methods

### Sample collection and isolation of plasma from pregnancy women

Plasma of pregnant women were acquired from collaborative hospitals. A written informed consent was obtained from the pregnant women while the blood was taken. Plasma was isolated by a continuous two-step centrifugation at 1600g for 10 minutes and 16000g for another 10 minutes, as described before [28]. The isolated plasma was stored at −80☐ for further use. All plasma samples were supplied with prenatal diagnosis results by amniocentesis and karyotyping and sanger sequencing, although the diagnostic information was not available to lab personnel while the study was conducted. Ethic approval was obtained from the Institutional Review Board of BGI.

### Isolation of exosomes from plasma of pregnancy women

Exosomes were isolated from plasma using the ExoQuick^TM^ Exosome Isolation Reagent (System Biosciences, America) following the manufacturer’s instructions. Briefly, 250μl plasma was treated with 2.5μl thrombin to a final concentration of 5U/mL for five minutes while mixing gently at room temperature. Centrifugation was then performed at 10,000rpm for fifteen minutes to remove visible fibrin pellet at the bottom. Supernatant was transferred to another Eppendorf tube and added with 63μl ExoQuick Exosome Precipitation Solution. The mixture was then cooled on ice for 30 minutes and centrifuged at 1500g for 30 minutes to remove all the trace of supernatant. The exosome pellet was finally resuspended in 100-200μl phosphate buffer saline (PBS) buffer. For flow cytometry analysis, the exosome pellet was resuspended in the Isolation Buffer (PBS with 0.1%BSA, filtered through a 0.22μm filter).

### Transmission electron microscopy

The isolated exosomes were diluted approximately 50 times with PBS and loaded on carbon-coated copper grids for 10 minutes. Excess liquid was removed by filter paper and then exosomes were negatively stained with 10ul of 2 % aqueous uranyl acetate solution for one minute. After dry at room temperature for 1.5 hour, exosomes were examined in a JEM-1230 transmission electron microscope (Japan Electronics Co., Ltd) at an accelerating voltage of 80 kV, magnification of 100,000 times to 200,000 times.

### Flow cytometry analysis

Exosomes were initially isolated using the ExoQuick^TM^ Exosome Isolation Reagent from 125μl plasma, as described above. Exosomes were then resuspended with 300ul Isolation Buffer. CD9 Dynabeads^®^ magnetic beads (Invitrogen, America) were added to the solution. The mixture was incubated overnight at 4☐ and another 60min at room temperature before separation to form exosome-beads complex. Then, a magnet was used to separate the complex from unbound exosomes. After washing twice with 200μl Isolation Buffer, the complex was resuspended with 200μl Isolation Buffer again and dispensed into two tubes. One tube was labeled as the sample tube, and 5μl antiCD63-Alexa647 antibody (#561983, BD Biosciences) was added. The other tube was labeled as the control tube, and 5μl IgG-Alexa647(#557714, BD Biosciences) was added. After rotationally incubating with the Alexa647-lablled antibody for 45 minutes at room temperature protected from light, the complex was separated from unbound antibody by the magnet and washed twice again with 200μl Isolation Buffer. Flow cytometry analysis was conducted using a FACSAria^TM^III (BD Biosciences) after resuspending the complex with 400μl Isolation Buffer.

### Western blotting

Exosome lysates were prepared by adding RIPA(P0013B, Beyotime, China), followed by centrifugation at 15,000g for 5 minutes. A total of 20μg exosome proteins determined by a Modified BCA Protein Assay Kit (Sangon Biotech, China) were loaded on a regular 8% SDS-PAGE gel. Electrophoresis was performed at 80V for 30 minutes followed at 120V for 90 minutes. Gels were transferred to polyvinylidene fluoride membranes (Millipore, Billerica) at 350mA for 70 minutes. The membranes were blocked with 5% skim milk overnight at 4☐. After incubation with primary antibodies at room temperature for three hours and secondary antibodies conjugated to horseradish peroxidase (HRP) at room temperature for one hour, the membranes were soaked with Pierce^TM^ ECL Western Blotting Substrate (Thermofisher, America), and chemiluminescence was detected using the instrument CLiNX. The following primary antibodies were used: rabbit anti-placenta alkaline phosphatase (PLAP) antibody (1:500; ABCAM), rabbit anti-CD63 antibody (1:1,000; ABCAM), mouse anti-calnexin antibody (1:500; Santa Cruz Biotechnology), mouse anti-CD81 antibody (1:500; Santa Cruz Biotechnology), mouse anti-CD9 antibody (1:500; Santa Cruz Biotechnology). The following secondary antibodies were used: goat anti-mouse IgG-HRP (1:4000; BPI), goat anti-rabbit IgG-HRP (1:4000; BPI). CD9, CD63, CD81 are typical exosome markers, while PLAP is a placenta specific marker, and Calnexin is an endoplasmic reticulum marker.

### ExoDNA and cfDNA extraction and fetal exoDNA detection

Before extracting exoDNA, exosomes were treated with two units of DNase I (New England Biolabs, America) for 1 hour to remove the residual cfDNA on the exosome surface. Then DNase I was then deactivated by 10mM EDTA. ExoDNA and cfDNA were separately extracted from the exosomes obtained from 250μl plasma using a Hipure Circulating DNA kit (Magen) following the manufacturer’s instructions. Qubit^TM^ dsDNA HS Assay kit(ThermoFisher, America) was used to confirm the quantity of cfDNA and exoDNA in 250μl plasma(Figure S1). The Quantifiler^®^ Trio DNA Quantification Kit (Applied Biosystems, America) were used to confirm the presence of DNA in exosomes using two pairs of primers targeting autosome, one pair of primers targeting Y chromosome and one pair of reference primers indicating the performance of PCR reaction. The △Ct difference value between cfDNA and exoDNA in equivalent plasma was calculated to reveal the relative quantity of exoDNA. The real-time quantitative-PCR (Q-PCR) reaction was performed with the ViiA 7 Real-Time PCR System (ABI).

### Massively parallel sequencing (MPS) of exoDNA and cfDNA

Pair-end (PE) sequencing libraries were constructed as described before [40], except that we increased the PCR cycle to 20 cycles. Briefly, end-repairing was performed at 37☐ for 10 minutes and adapter ligation of barcodes was performed at 23☐ for 30 minutes. Ligation products were then purified with Agencourt AMPure XP beads (BECKMAN COULTER), followed by 20 cycles of PCR amplification and further purification. The PCR products were heat-denatured at 95☐ and cooled on ice for three minutes to generate single strand DNA for ligation using T4 ligase. PE sequencing was conducted with the BGISEQ-500 sequencer (MGI, China). 2 x 50bp PE sequencing with 30M data was obtained for each sample.

### Multiplex PCR

Anchored multiplex PCR(AMP) of exoDNA was conducted to amplify the coding region of the *FGFR3* gene (OMIM:134934). exoDNA was extracted after exosome digestion step and then the target amplicon library was constructed by AMP technology [41]. Briefly, exoDNA is processed with end repair and dA tailing, directly followed by ligation with adapter containing barcodes. Solid-phase reversible immobilization(SPRI)-cleaned by Agencourt AMPure XP beads, ligated fragments are amplified with 20 cycles of multiplex PCR1 using gene-specific primers(Table S3 Upstream primers). SPRI-cleaned PCR1 amplicons are amplified with a second round of 20 cycles multiplex PCR2(Table S3 Downstream primers). After a final SPRI cleanup, the target amplicon library is ready for quantitation and sequencing. Sequencing performed on BGISEQ-500RS.

### Bioinformatics

Adaptors and reads with low quality were removed. Then, PE sequencing reads were mapped to the human reference genome (Hg19, GRCh37) using the BWA software and the insert size of cfDNA and exoDNA was calculated according to the bam file. Then we calculated the GC content and relative reads ratio in every 1M window size of all the chromosomes, and visualized it with Circos package in R. The coefficient of variation (CVmean) of each sample is the mean CV of the 22 autosomes:

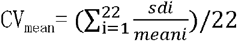

mean_i_ and sd_i_ are the mean reads number and sd of all the windows in each chromosome separatively.

The mitochondrial DNA percent can correspond to the proportion of reads mapped uniquely to the mitochondrial DNA genome and nuclear genome.

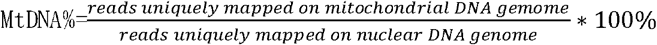

Fetal fraction was deduced with the unique reads mapped to the chromosome Y, and the method to calculate fetal fraction was described previously [29].

Fetal risk of chromosome aneuploid were calculated using the similar strategy to cfDNA NIPT [28,29]. Generally, the disease risk was detected by a binary hypothesis t-test that calculating the z-score of the risk. Using a cutoff value of 3, z-score > 3 means the high risk of disease. Fetal risk of monogenic diseases was evaluated by evaluating the minor allele frequency (MAF), which was calculated by the ratio between the reads of the second most common allele and the total reads of the SNP location detected.

R package (R-3.2.5) were used to carry out the t-test and Pearson correlation analysis.

## Acknowledgments

This work was supported by the National Natural Science Foundation of China(No.81601294), Shenzhen Birth Defects Screening Project Lab (JZF No. [2016]750), Shenzhen Municipal Government of China (NO. JCYJ20150403101146312) and Guangzhou Science and Technology Program (No. 201604020078).

## Author Contributions

All authors confirmed they have contributed to the intellectual content of this paper and have met the following 3 requirements: (a) significant contributions to the conception and design, acquisition of data, or analysis and interpretation of data; (b) drafting or revising the article for intellectual content; and (c) final approval of the published article.

## Supplementary Materials

**Table 1** Clinical information of the 20 euploidy, 9 T21, 3 T18 and 1 T13 plasma of pregnancy women.

**Table 2** Clinical information of the achondroplasia and thanatophoric dysplasia.

**Table 3** Primers target *FGFR3* in AMP method.

**Fig.S1.**
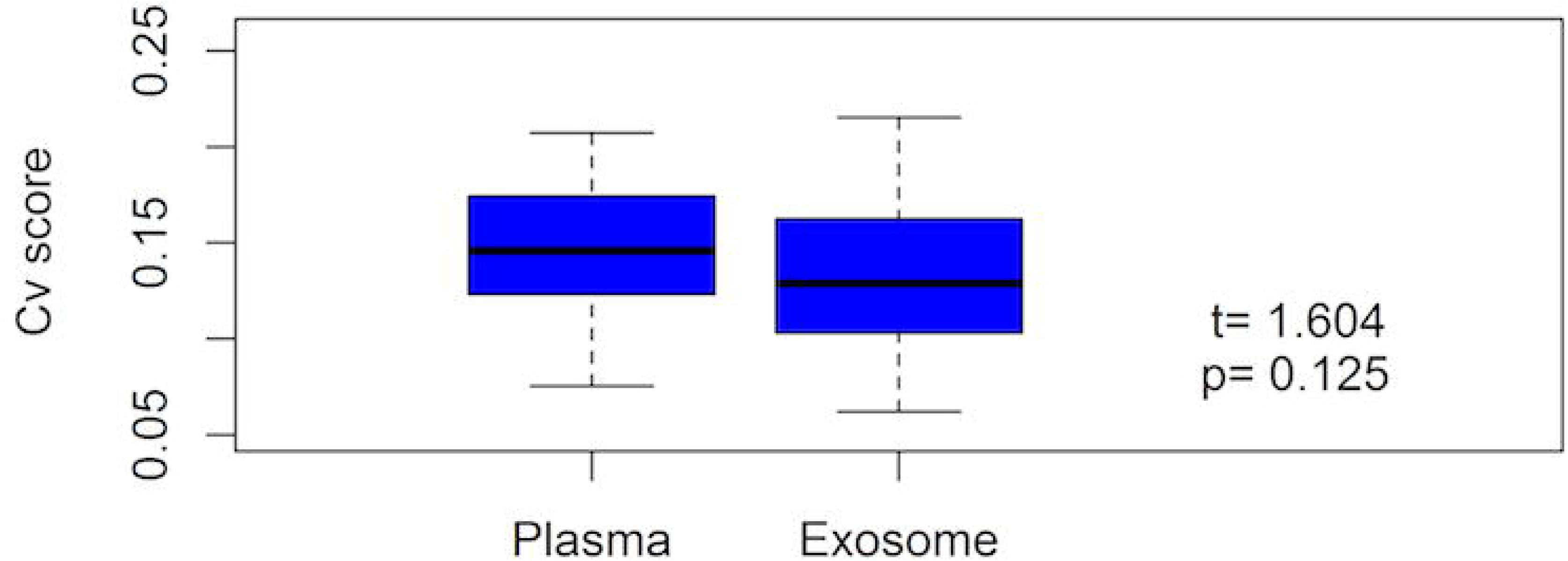
Qubit result of the extracted exoDNA and cfDNA in 20μl elution buffer from 250μl plasma of 20 euploidy.

**Fig.S2.**
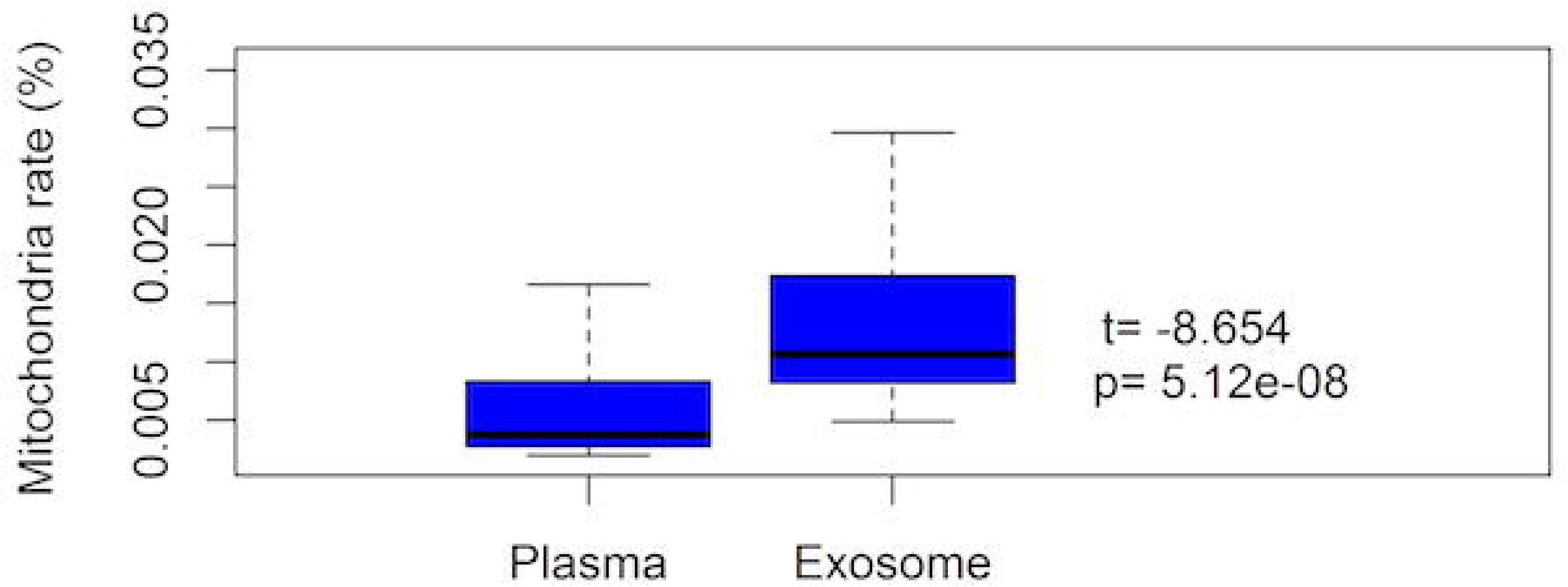
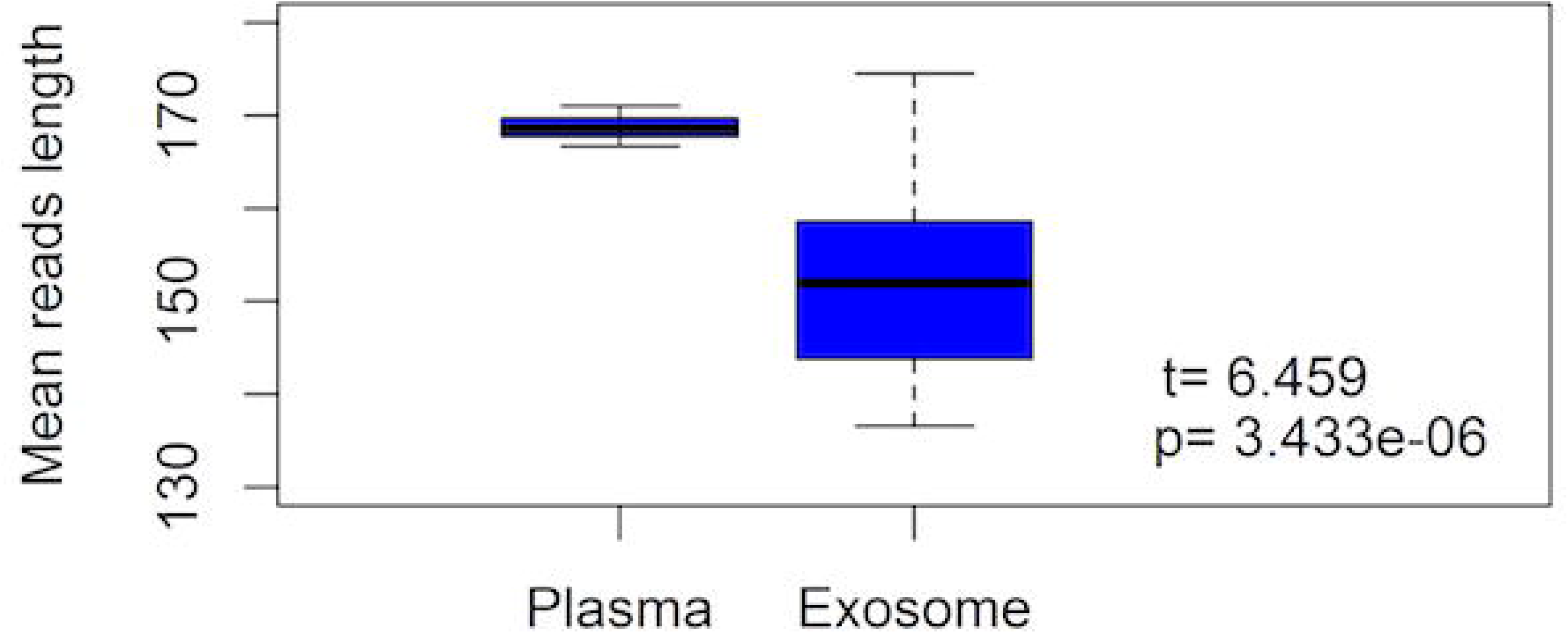
The fragment size distribution of exoDNA(blue) and cfDNA(red) from 20 euploidy.

**Fig.S3.**
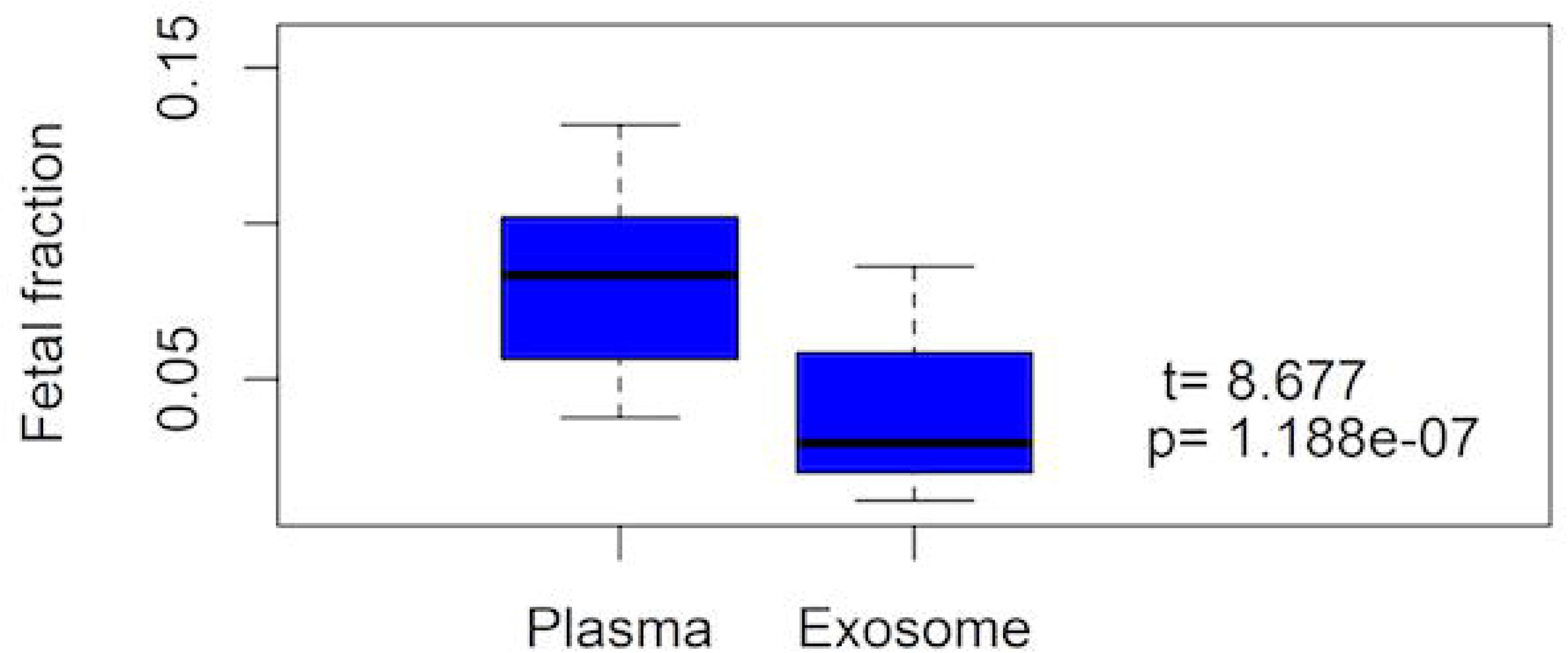
Regression analysis of fetal fraction between cfDNA and exoDNA, and fetal fraction of exoDNA and gestational days.

## References

1. Colombo M, Raposo G, Thery C (2014) Biogenesis, secretion, and intercellular interactions of exosomes and other extracellular vesicles. Annu Rev Cell Dev Biol 30: 255–289.

2. Yanez-Mo M, Siljander PR, Andreu Z, Zavec AB, Borras FE, et al. (2015) Biological properties of extracellular vesicles and their physiological functions. J Extracell Vesicles 4: 27066.

3. Cai J, Wu G, Jose PA, Zeng C (2016) Functional transferred DNA within extracellular vesicles. Exp Cell Res 349: 179–183.

4. Cai J, Han Y, Ren H, Chen C, He D, et al. (2013) Extracellular vesicle-mediated transfer of donor genomic DNA to recipient cells is a novel mechanism for genetic influence between cells. J Mol Cell Biol 5: 227–238.

5. Sarker S, Scholz-Romero K, Perez A, Illanes SE, Mitchell MD, et al. (2014) Placenta-derived exosomes continuously increase in maternal circulation over the first trimester of pregnancy. J Transl Med 12: 204.

6. Salomon C, Torres MJ, Kobayashi M, Scholz-Romero K, Sobrevia L, et al. (2014) A gestational profile of placental exosomes in maternal plasma and their effects on endothelial cell migration. PLoS One 9: e98667.

7. Mitchell MD, Peiris HN, Kobayashi M, Koh YQ, Duncombe G, et al. (2015) Placental exosomes in normal and complicated pregnancy. American journal of obstetrics and gynecology 213: S173–S181.

8. Mincheva-Nilsson L, Baranov V (2014) Placenta-derived exosomes and syncytiotrophoblast microparticles and their role in human reproduction: immune modulation for pregnancy success. Am J Reprod Immunol 72: 440–457.

9. Salomon C, Scholz-Romero K, Sarker S, Sweeney E, Kobayashi M, et al. (2016) Gestational Diabetes Mellitus Is Associated With Changes in the Concentration and Bioactivity of Placenta-Derived Exosomes in Maternal Circulation Across Gestation. Diabetes 65: 598–609.

10. Luo SS, Ishibashi O, Ishikawa G, Ishikawa T, Katayama A, et al. (2009) Human villous trophoblasts express and secrete placenta-specific microRNAs into maternal circulation via exosomes. Biol Reprod 81: 717–729.

11. Baig S, Lim J, Fernandis A, Wenk M, Kale A, et al. (2013) Lipidomic analysis of human placental syncytiotrophoblast microvesicles in adverse pregnancy outcomes. Placenta 34: 436–442.

12. Baig S, Kothandaraman N, Manikandan J, Rong L, Ee KH, et al. (2014) Proteomic analysis of human placental syncytiotrophoblast microvesicles in preeclampsia. Clinical proteomics 11: 40.

13. Kahlert C, Melo SA, Protopopov A, Tang J, Seth S, et al. (2014) Identification of double-stranded genomic DNA spanning all chromosomes with mutated KRAS and p53 DNA in the serum exosomes of patients with pancreatic cancer. J Biol Chem 289: 3869–3875.

14. Thakur BK, Zhang H, Becker A, Matei I, Huang Y, et al. (2014) Double-stranded DNA in exosomes: a novel biomarker in cancer detection. Cell Res 24: 766–769.

15. Lazaro-Ibanez E, Sanz-Garcia A, Visakorpi T, Escobedo-Lucea C, Siljander P, et al. (2014) Different gDNA content in the subpopulations of prostate cancer extracellular vesicles: apoptotic bodies, microvesicles, and exosomes. Prostate 74: 1379–1390.

16. San Lucas FA, Allenson K, Bernard V, Castillo J, Kim DU, et al. (2016) Minimally invasive genomic and transcriptomic profiling of visceral cancers by next-generation sequencing of circulating exosomes. Ann Oncol 27: 635–641.

17. Guescini M, Genedani S, Stocchi V, Agnati LF (2010) Astrocytes and Glioblastoma cells release exosomes carrying mtDNA. J Neural Transm (Vienna) 117: 1–4.

18. Lo YD, Chan KA, Sun H, Chen EZ, Jiang P, et al. (2010) Maternal plasma DNA sequencing reveals the genome-wide genetic and mutational profile of the fetus. Science translational medicine 2: 61ra91–61ra91.

19. Hudecova I, Sahota D, Heung MM, Jin Y, Lee WS, et al. (2014) Maternal plasma fetal DNA fractions in pregnancies with low and high risks for fetal chromosomal aneuploidies. PloS one 9: e88484.

20. Zhang H, Gao Y, Jiang F, Fu M, Yuan Y, et al. (2015) Non‐invasive prenatal testing for trisomies 21, 18 and 13: clinical experience from 146 958 pregnancies. Ultrasound in Obstetrics & Gynecology 45: 530–538.

21. Chen Y, Zhao L, Wang Y, Cao M, Gelowani V, et al. (2017) SeqCNV: a novel method for identification of copy number variations in targeted next-generation sequencing data. BMC bioinformatics 18: 147.

22. Chen S, Lau TK, Zhang C, Xu C, Xu Z, et al. (2013) A method for noninvasive detection of fetal large deletions/duplications by low coverage massively parallel sequencing. Prenatal diagnosis 33: 584–590.

23. Salomon C, Kobayashi M, Ashman K, Sobrevia L, Mitchell M, et al. (2013) Hypoxia-induced changes in the bioactivity of cytotrophoblast-derived exosomes. PLoS ONE 8: e79636.

24. Mincheva-Nilsson L, Baranov V (2010) The role of placental exosomes in reproduction. American journal of reproductive immunology 63: 520–533.

25. Dragovic R, Collett G, Hole P, Ferguson D, Redman C, et al. (2015) Isolation of syncytiotrophoblast microvesicles and exosomes and their characterisation by multicolour flow cytometry and fluorescence Nanoparticle Tracking Analysis. Methods 87: 64–74.

26. Bischoff FZ, Lewis DE, Simpson JL (2005) Cell-free fetal DNA in maternal blood: kinetics, source and structure. Hum Reprod Update 11: 59–67.

27. Fernando M, Jiang C, Krzyzanowski G, Ryan W (2017) New evidence that a large proportion of human blood plasma cell-free DNA is localized in exosomes. PLoS ONE 12: e0183915.

28. Zhang H, Gao Y, Jiang F, Fu M, Yuan Y, et al. (2015) Non-invasive prenatal testing for trisomies 21, 18 and 13: clinical experience from 146,958 pregnancies. Ultrasound Obstet Gynecol 45: 530–538.

29. Jiang F, Ren J, Chen F, Zhou Y, Xie J, et al. (2012) Noninvasive Fetal Trisomy (NIFTY) test: an advanced noninvasive prenatal diagnosis methodology for fetal autosomal and sex chromosomal aneuploidies. BMC Med Genomics 5: 57.

30. Mustapic M, Eitan E, Werner J, Berkowitz S, Lazaropoulos M, et al. (2017) Plasma Extracellular Vesicles Enriched for Neuronal Origin: A Potential Window into Brain Pathologic Processes. Front Neurosci 11: 278.

31. Hong CS, Muller L, Boyiadzis M, Whiteside TL (2014) Isolation and characterization of CD34+ blast-derived exosomes in acute myeloid leukemia. PLoS One 9: e103310.

32. He M, Crow J, Roth M, Zeng Y, Godwin AK (2014) Integrated immunoisolation and protein analysis of circulating exosomes using microfluidic technology. Lab Chip 14: 3773–3780.

33. Reategui E, van der Vos KE, Lai CP, Zeinali M, Atai NA, et al. (2018) Engineered nanointerfaces for microfluidic isolation and molecular profiling of tumor-specific extracellular vesicles. Nat Commun 9: 175.

34. Lv L-L, Cao Y, Liu D, Xu M, Liu H, et al. (2013) Isolation and Quantification of MicroRNAs from Urinary Exosomes/Microvesicles for Biomarker Discovery. International Journal of Biological Sciences 9: 1021–1031.

35. Mall C, Rocke D, Durbin-Johnson B, Weiss R (2013) Stability of miRNA in human urine supports its biomarker potential. Biomark Med 7: 623–631.

36. Ge Q, Zhou Y, Lu J, Bai Y, Xie X, et al. (2014) miRNA in plasma exosome is stable under different storage conditions. Molecules 19: 1568–1575.

37. Sabapatha A, Gercel-Taylor C, Taylor D (2006) Specific isolation of placenta-derived exosomes from the circulation of pregnant women and their immunoregulatory consequences. Am J Reprod Immunol 56: 345–355.

38. Menon R, Mesiano S, Taylor RN (2017) Programmed Fetal Membrane Senescence and Exosome-Mediated Signaling: A Mechanism Associated With Timing of Human Parturition. Front Endocrinol (Lausanne) 8: 196.

39. Takahashi A, Okada R, Nagao K, Kawamata Y, Hanyu A, et al. (2017) Exosomes maintain cellular homeostasis by excreting harmful DNA from cells. Nat Commun 8: 15287.

40. Huang J, Liang X, Xuan Y, Geng C, Li Y, et al. (2017) A reference human genome dataset of the BGISEQ-500 sequencer. Gigascience 6: 1–9.

41. Zheng Z, Liebers M, Zhelyazkova B, Cao Y, Panditi D, et al. (2014) Anchored multiplex PCR for targeted next-generation sequencing. Nature medicine 20: 1479.

